# Towards Decoding the Metabolic Plasticity in Cancer: Coupling of Gene Regulation and Metabolic Pathways

**DOI:** 10.1101/428367

**Authors:** Dongya Jia, Mingyang Lu, Kwang Hwa Jung, Jun Hyoung Park, Linglin Yu, José N. Onuchic, Benny Abraham Kaipparettu, Herbert Levine

**Author notes:** Correspondence: Herbert Levine, Benny Abraham Kaipparettu, José N. Onuchic.

## Abstract

Metabolic plasticity enables cancer cells to switch their metabolism phenotypes between glycolysis and oxidative phosphorylation (OXPHOS) during tumorigenesis and metastasis. However, it is still largely unknown how cancer cells orchestrate gene regulation to balance their glycolysis and OXPHOS activities for better survival. Here, we establish a theoretical framework to model the coupling of gene regulation and metabolic pathways in cancer. Our modeling results demonstrate a direct association between the activities of AMPK and HIF-1, master regulators of OXPHOS and glycolysis respectively, with the activities of three metabolic pathways: glucose oxidation, glycolysis and fatty acid oxidation (FAO). Guided by the model, we develop metabolic pathway signatures to quantify the activities of glycolysis, FAO and the citric acid cycle of tumor samples by evaluating the expression levels of enzymes involved in corresponding processes. The association of AMPK/HIF-1 activity with metabolic pathway activity, predicted by the model and verified by analyzing the gene expression and metabolite abundance data of patient samples, is further validated by in vitro studies of aggressive triple negative breast cancer cell lines.

## Introduction

Abnormal metabolism is an emerging hallmark of cancer (1, 2). Unlike normal cells, cancer cells largely depend on glycolysis to produce energy even in presence of oxygen, referred to as the Warburg effect (3) or aerobic glycolysis. The upregulation of glucose transporters (GLUTs) (4) and glycolytic enzymes, such as lactate dehydrogenase (LDH) and hexokinase 2 (HK2), has been well documented in multiple types of cancer cells (2, 5). Cancer cells can use aerobic glycolysis for rapid ATP production and biomass synthesis to facilitate tumorigenesis and modulate their metastatic potential (6). Moreover, the enhanced glycolytic activity of cancer cells has been shown to be associated with increased therapy resistance [7–9].

It has been becoming clear that in addition to aerobic glycolysis, oxidative phosphorylation (OXPHOS) can also plays critical roles in various types of cancer [9–12]. For example, the tumor progression and metastatic propensity of several triple negative breast cancer (TNBC) cell lines and patient-derived xenograft (PDX) models are largely affected by their energy dependency on FAO. Pharmacologic repression of FAO activity significantly inhibits their *in vivo* tumor growth (12, 14). The mouse basal breast cancer cell line 4T1, which is part of an isogenic model system, is supermetastatic and commonly used to simulate the human stage IV breast cancer (15). Compared to its isogenic non-metastatic 67NR cells, 4T1 cells exhibit both enhanced OXPHOS and increased glycolytic activities (16). Moreover, the circulating tumor cells (CTCs) derived from 4T1 cells show significantly higher mitochondrial respiration and biogenesis activities as compared to both its primary tumors and its lung metastases (17). Notably, there is no observable decrease in glycolytic activity of these 4T1 CTCs, indicating the co-existence of OXPHOS and glycolysis. The association of high OXPHOS activity with high metastatic potential has also been observed in mouse melanoma B16-M4b cells and human cervical cancer SiHa-F3 cells [9]. All these indicate that cancer cells are able to utilize both glycolysis and OXPHOS, depending on its circumstances. However, it is still largely unknown how cancer cells orchestrate the metabolic pathway activity through gene regulation to facilitate malignancy.

Mathematical modeling approaches have been employed to help elucidate the aforementioned metabolic reprogramming in cancer. Constraint-based models including flux balance analysis (FBA) based on conservation of mass (18, 19) have been the most widely used approaches to simulate cancer metabolism (20, 21). Applications include quantifying the extent of aerobic glycolysis of the NCI-60 cell lines (22), characterizing their primary uptake pathways (23) and identifying the change of their metabolic states upon gene knockdown and/or drug treatment (24).In addition to the characterization of the metabolic flux change in cancer cells, modeling efforts have also been developed to identify anomalous activity for genes that are involved in controlling cancer metabolism. For example, there are studies modeling the effects of reactive oxygen species (ROS) and antioxidants on hypoxia-inducible factor 1 (HIF-1), a master regulator of glycolysis (25), and modeling the interplay between HIF-1 and 5’ AMP-activated protein kinase (AMPK), a master regulator of OXPHOS and mitochondrial biogenesis (26). These computational works offer a quantitative and dynamical perspective of cancer metabolism mostly focusing on either metabolic pathway activity or gene activity. However, the alteration of the metabolic activity is coupled with the change in gene activity, and *vice versa*. For example, HIF-1, that is often stabilized in cancer cells due to hypoxia, promotes glycolytic activity by upregulating the expression of glucose transporter genes, such as GLUT1, and glycolytic enzyme genes, such as pyruvate kinase isozyme M2 (PKM2) (27). Upregulation of PKM2 in turn promotes HIF-1 transactivation and the products of glycolysis, such as lactate, further stabilizes HIF-1, thus forming self-enforcing feedback loops of HIF-1 activity (28). Thus, to comprehensively characterize cancer metabolic reprogramming, a mathematical modeling framework integrating gene regulation with metabolic pathways is urgently needed.

Previously, to decipher the genetic interplay between glycolysis and OXPHOS, we developed a mathematical model focusing on a core regulatory circuit, composed of AMPK, HIF-1 and ROS (26). Computational modeling of this circuit showed that cancer cells can robustly acquire three stable steady states - ‘W’ (HIF-1^high^/pAMPK^low^), ‘O’ (HIF-1^low^/pAMPK^high^) and ‘W/O’ (HIF-1^high^/ pAMPK^high^) - corresponding to a glycolysis phenotype, an OXPHOS phenotype and a hybrid metabolic phenotype, in which cancer cells use both glycolysis and OXPHOS. Here, the AMPK activity is represented by the level of phosphorylated AMPK (pAMPK) at threonine-172 of the α subunit. The emergence of the hybrid metabolic phenotype of cancer cells from our modeling analysis has been supported by recent experimental evidence (10, 12, 13, 16, 17). We argued that the hybrid metabolic phenotype exhibits metabolic plasticity to adapt to varying microenvironments and hence tends to be more aggressive relative to cells in a more glycolysis or OXPHOS phenotype (13).

Here, to capture the coupling of gene activity and metabolic pathway activity, we extend our AMPK:HIF-1:ROS model by coupling it to three distinct metabolic pathways, glycolysis, glucose oxidation and FAO. Unlike traditional FBA, our model captures gene regulation and pathway activity and their interactions via chemical rate equations. The extended model elucidates the detailed association of AMPK and HIF-1 activities with metabolic pathway activities for each stable state. High AMPK activity associates with high OXPHOS pathway activity (glucose oxidation and/or FAO) and high HIF-1 activity associates with high glycolysis activity. Particularly, the hybrid metabolic state ‘W/O’ is characterized by HIF-1^high^/pAMPK^high^ and high activity of both glycolysis and OXPHOS (glucose oxidation and/or FAO). Our predicted association of AMPK/HIF-1 activity with metabolic pathway activity is confirmed by analyzing well-annotated metabolomic and transcriptomic data from a breast cancer patients’ cohort. This utilizes metabolic pathway activity signatures developed in the present work as well as our published AMPK and HIF-1 signatures (26). This result was further validated in human invasive breast carcinoma and hepatocellular carcinoma (HCC) using RNA-seq data from The Cancer Genome Atlas (TCGA). For direct experimental validation, we use two TNBC cell lines, SUM159-PT and MDA-MB-231 and confirm that metastatic TNBC cells can acquire a stable hybrid metabolic phenotype. We further show that repressing the glycolytic activity activates AMPK and enhances OXPHOS activity, and conversely repressing mitochondrial function upregulates multiple glycolysis genes and increases glycolytic activity. Finally, a combination of glycolytic and OXPHOS inhibitors that potentially eliminates the existence of the hybrid metabolic phenotype exhibits maximum reduction of proliferation and clonogenicity of these TNBC cells. In summary, through integrating mathematical modeling, bioinformatics and experiments, we demonstrate a direct association of the AMPK/HIF-1 activity with metabolic pathway activity and investigate the existence of the aggressive hybrid metabolic phenotype.

## Results

### The regulatory network of glycolysis and OXPHOS coupling gene regulation and metabolic pathways

Our starting point is a diagram constructed from an extensive literature survey summarizing important gene regulation and energy pathways (**Fig. 1**). AMPK, the mitochondrial energy sensor, can stimulate glucose and lipid oxidation for energy production in response to metabolic stress. Specifically, AMPK can promote glucose uptake through GLUT4 translocation and enhance the OXPHOS activity and mitochondrial biogenesis by activating the transcription factors peroxisome proliferator-activated receptor gamma coactivator 1-alpha (PGC-1α) and cAMP response element-binding protein (CREB). AMPK can also activate FAO by increasing the activity of carnitine palmitoyltransferase-1 (CPT-1) via repressing acetyl-CoA carboxylase-2 (ACC2) through phosphorylation (29). HIF-1 is often stabilized in cancer and can promote glycolytic activity by upregulating the expression of glucose transporters and glycolytic enzymes (27) and the products of glycolysis, such as lactate, can in turn stabilize HIF-1 (28). HIF-1 can also stimulate the uptake of glucose to fuel glycolysis (30). Notably, there are mutually inhibitory feedback loops between AMPK and HIF-1. HIF-1 directly represses the transcription of AMPK and AMPK inhibits HIF-1 via the mTOR signaling pathway (31).

**Figure 1.**
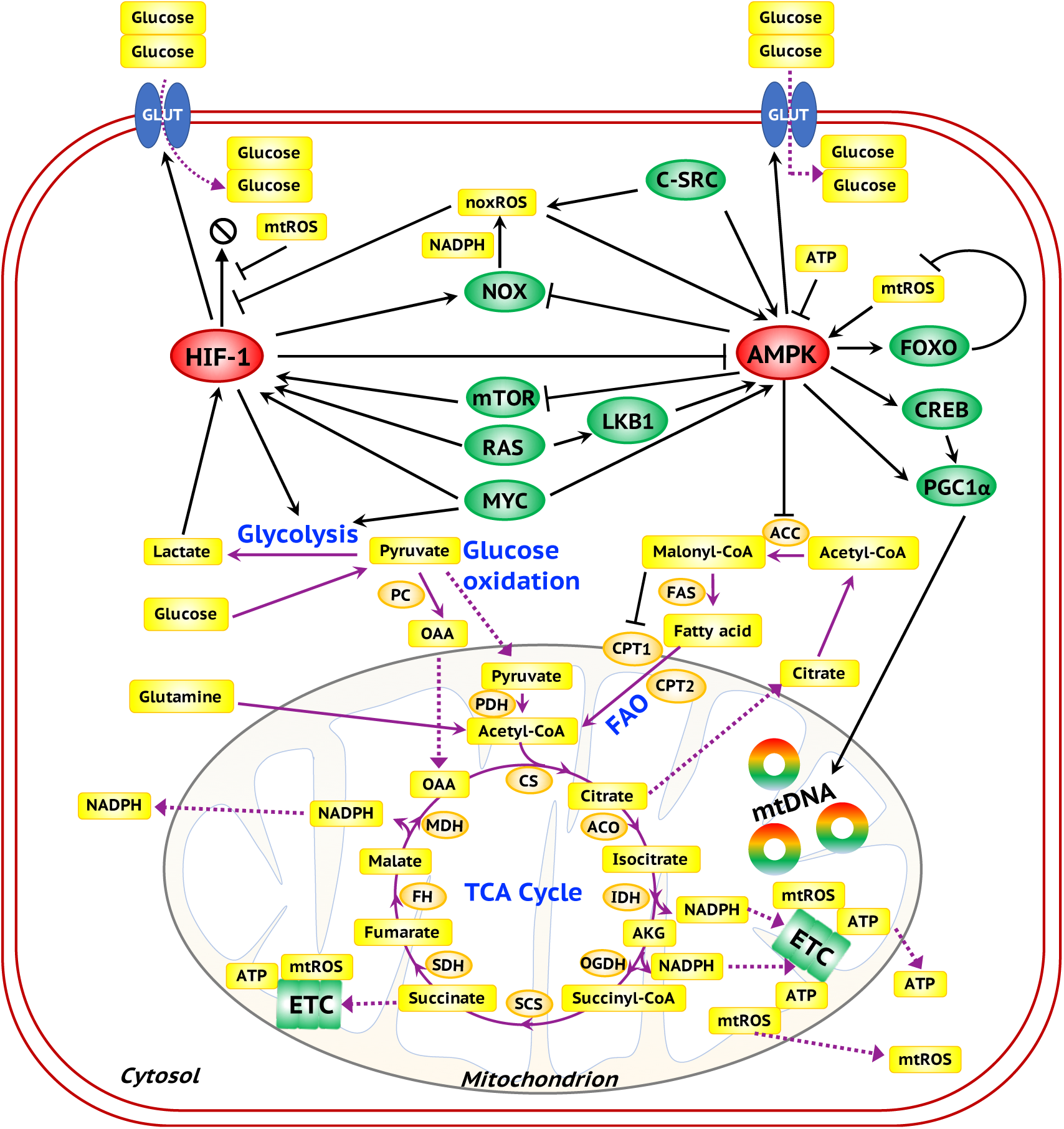
A comprehensive regulatory network of glycolysis and OXPHOS. The ovals represent genes. Red ovals highlight the master regulators AMPK and HIF-1. Green ovals represent downstream target genes of the master regulators and oncogenes. Orange ovals represent the enzyme genes. The yellow rectangles represent metabolites. The black arrows represent excitatory regulation and the black bar-headed arrows represent inhibitory regulation. The purple solid lines represent the chemical reactions in metabolic pathways and the purple dotted lines represent the transportation of metabolites.

Intracellular glucose is a carbohydrate resource for glycolysis and glucose oxidation. Both glucose oxidation and FAO can generate acetyl-CoA to fuel the tricarboxylic acid cycle (TCA cycle) and subsequently generate ATP via the electron transport chain (ETC). The capacity of mitochondria to utilize acetyl-CoA to fuel TCA is regulated by AMPK. ROS, including mitochondrial ROS (mtROS) generated during OXPHOS (glucose oxidation and FAO) and the NOX-derived ROS (noxROS) produced in the cytosol, can stabilize HIF-1 and activate AMPK [28–31]. Activation of AMPK can in turn trigger a deoxidation program through upregulating the transcription factor forkhead box O (FOXO) to increase the expression of thioredoxin to avoid excessive ROS caused damage [32–34]. All three metabolic pathways considered here, glucose oxidation, glycolysis and FAO, generate ATP and excessive ATP can repress the activity of AMPK to avoid its over-activation (32). In addition, many oncogenes, such as c-Src, RAS, MYC, are actively involved in the regulation of cancer metabolism through modulating the activity of AMPK, HIF-1 and enzymes in multiple metabolic pathways. Note that some aspects of metabolism such as the glutamine pathway or the relevance of autophagy are beyond the scope of this paper and will be treated in future extensions of this research.

### Coupling the AMPK:HIF-1:ROS circuit with glycolysis/OXPHOS pathways

To create a computational model which can capture the basic principles of the coupling between gene regulation and metabolic pathways, we coarse-grain the extensive regulatory system (**Fig. 1**) into a minimum network consisting of the AMPK:HIF-1:ROS core regulatory circuit, the three metabolic pathways, glycolysis, glucose oxidation and FAO (**Fig. 2**). AMPK and HIF-1 form mutually inhibitory feedback loops and AMPK reduces while HIF-1 increases noxROS which in turn activates both AMPK and HIF-1 (26). We now extend this AMPK:HIF-1:ROS model by explicitly including the metabolic pathways. The effective self-activation of HIF-1 is through its interaction with the glycolytic pathway and the effective self-inhibition of AMPK is due to its interaction with OXPHOS processes. One critical component of our modeling is the direct consideration of glucose levels and acetyl-CoA levels. Glucose is the competing resources for glycolysis and glucose oxidation and acetyl-CoA, as the only fuel entering TCA cycle, is the common intermediate of both glucose oxidation and FAO. The uptake rate of glucose is regulated by AMPK and HIF-1 and the utilization rate of acetyl-CoA for TCA cycle is restricted by the mitochondrial capacity, which is regulated by AMPK. The detailed formulation of the mathematical model representing the dynamics of pAMPK, HIF-1, mtROS, noxROS, ATP, glucose, acetyl-CoA together with the rates of glycolysis, glucose oxidation and FAO is provided in the ‘**Methods and Materials**’ section. To reconcile the different time scales of gene regulation and metabolic flux, we always assume that the metabolite concentration and the pathway activities are in a steady state at certain levels of pAMPK and HIF-1 since metabolic processes are much faster than gene regulation. We propose that our coarse-grained network captures the key features of the more comprehensive set of interactions indicated in **Fig. 1** and in particular is sufficient to explain important experimental observations on the coupling of gene activity and metabolic pathway activity.

**Figure 2.**
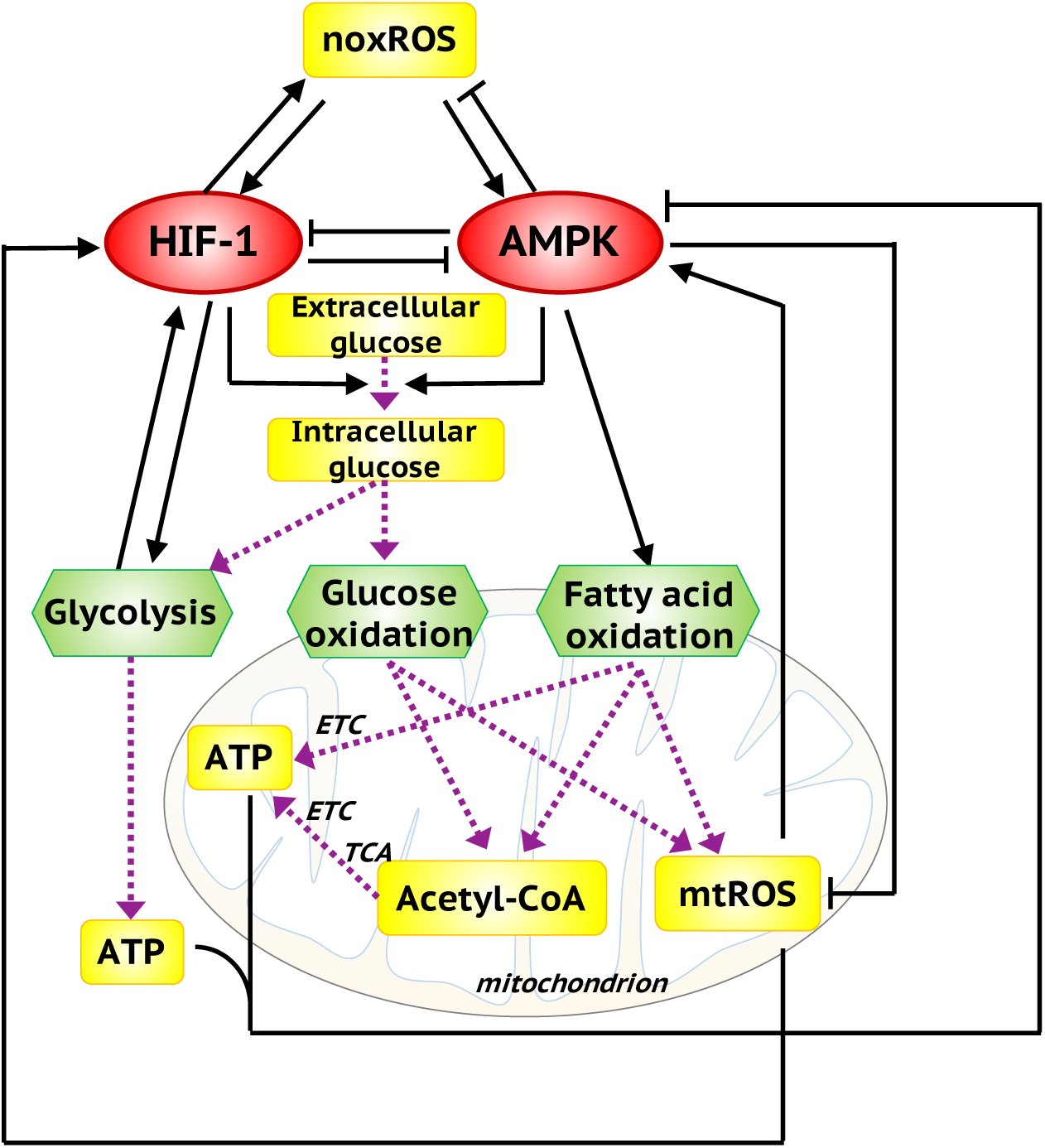
The metabolic regulatory network, coupling the core AMPK:HIF-1:ROS circuit with three metabolic pathways. AMPK and HIF-1 are the master regulators of OXPHOS and glycolysis. Both AMPK and HIF-1 can promote glucose uptake. The intracellular glucose can be used by glycolysis and glucose oxidation. Both glucose oxidation and FAO can generate acetyl-CoA to fuel the TCA cycle and consequently the production of ATP via ETC. mtROS and ATP can in turn regulate the activity of AMPK and HIF-1. Here, the black solid arrows/bar-headed arrows represent regulatory links. The purple dotted arrows represent the metabolic pathways.

### Genetic and metabolic characterization of each metabolism phenotype

First, we analyze the mathematical equations that were derived to simulate the coarse-grained network using two sets of parameters corresponding to normal and cancer cells respectively. Cancer cells often have higher mtROS production rate due to reprogrammed mitochondria (10) and more stabilized HIF-1 due to the typically hypoxic conditions. Therefore, the value of the parameters 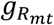 representing the production rates of mtROS during OXPHOS (glucose oxidation and FAO) is taken to be larger, and the value of the parameter *k_H_* representing the degradation rate of HIF-1 smaller for cancer relative to normal cells. The values of the other parameters are unchanged. We will consider in more detail the effects of mtROS production rate and HIF-1 degradation rate on cancer metabolic plasticity in a later section.

To identify the robust stable metabolic states enabled by the regulatory network (**Fig. 2**), we utilize a parameter randomization approach. The overall strategy involves randomizing the modeling parameters for each simulation and collecting all stable steady solutions for statistical analysis, by which the most significant solution patterns can be identified (33, 34). As expected, the solution patterns are conserved even in the presence of large parameter perturbations due to restraints from the network topology, i.e. extensive crosstalk of regulatory proteins and energy pathways. We consider 1000 sets of model parameters and for each set, the value of each parameter, except for the fixed values of mtROS production rate and HIF-1 degradation rate that distinguish cancer cells from normal cells, is randomly sampled from (75%*p*_0_, 125%*p*_0_), where *p*_0_ is the baseline value. We collect all of the stable state solutions and use unsupervised hierarchical clustering analysis (HCA) to identify the patterns present in the solution set. HCA shows that the stable state solutions form three large clusters; one is characterized by high pAMPK/mtROS/G1/F, low HIF-1/noxROS/G2 (G1 represents the glucose oxidation rate, F represents the FAO rate and G2 represents the glycolysis rate), corresponding to an OXPHOS state, one is characterized by high HIF-1/noxROS/G_2_, low pAMPK/mtROS/G1/F, corresponding to a glycolytic state, and one is characterized by high pAMPK/mtROS/G1/F, high HIF-1/noxROS/G2, corresponding to a hybrid metabolic state (**Fig. 3A, Fig S1**).

**Figure 3.**
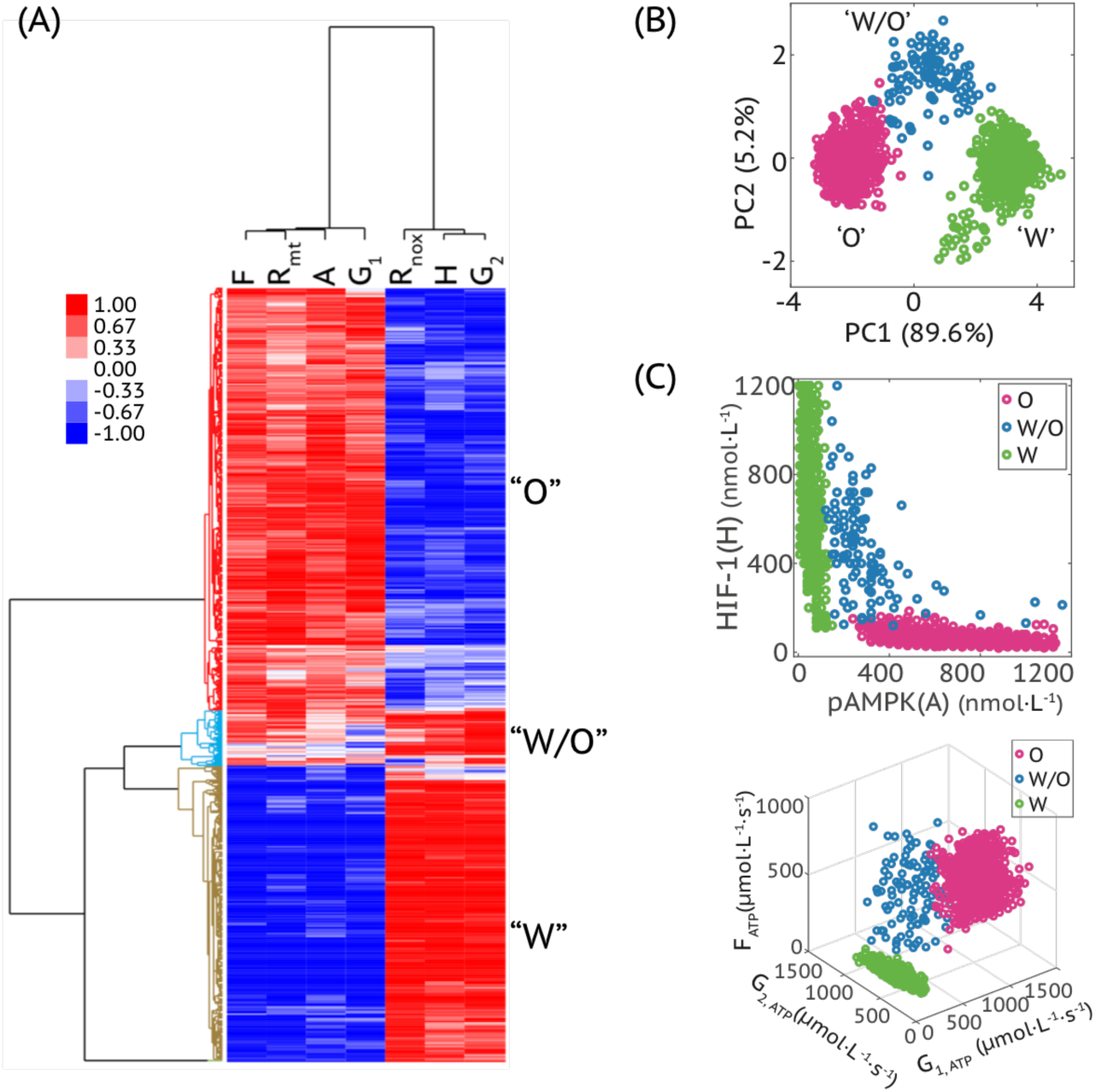
Modeling prediction of the association between AMPK/HIF-1 activity and metabolic pathway activity. (A) Hierarchical clustering analysis of the stable state solutions from 1000 sets of parameters. Each row represents one stable state solution, referred to as one sample here and each column represents the levels of a regulatory protein, metabolite or the rates of one metabolic pathway. In (A), F represents FAO rate; R_mt_ represents mtROS level; A represents pAMPK level; G_1_ represents glucose oxidation rate; R_nox_ represents noxROS level; H represents HIF-1 level; G_2_ represents glycolysis rate. The solutions can be clustered into three main groups, referred to as cluster “O”, “W/O” and “W”, as marked by different colors in the dendrogram. (B) Principal component analysis of the clustered samples in (A). (C) Top panel: The pAMPK and HIF-1 levels of the clustered samples in (A). Bottom panel, the metabolic pathway activities of the clustered samples in (A). The colors representing different clusters - “O”, “W/O” and “O” - are consistent with those used in (B).

We also visualized the three clusters by projecting the stable state solutions onto the first and second principal components (PC1 and PC2) through principal component analysis (PCA) of all solutions (**Fig. 3B**), onto the pAMPK and HIF-1 axes and onto the G1, G2 and F axes (**Fig. 3C**). These results indicate that cells in the ‘W’ state mostly use glycolysis for ATP production, cells in the ‘O’ state mainly use OXPHOS (including glucose oxidation and FAO) for ATP production, while cells in the hybrid ‘W/O’ state can utilize all three metabolic pathways to generate ATP (**Fig. 3C**). Thus, the modeling analysis demonstrates an association of high AMPK activity with high OXPHOS activity, and high HIF-1 activity with high glycolytic activity. This was conjectured to be the case in our previous work which however did not include any explicit analysis of the metabolic processes. Similarly, we performed an analogous analysis for the normal cells. However, the hybrid metabolic state is rarely observed among the stable state solutions from 1000 sets of randomized parameters (**Fig. S2**).

### The effects of HIF-1 degradation rate and mtROS production rate in modulating cancer metabolic phenotypes

The modeling framework can be utilized to analyze the effects of various kinds of perturbations on cancer metabolic phenotypes. In this section, using the HIF-1 degradation rate and the mtROS production rate as two examples, we will show how changing these two variables would modify cancer metabolism phenotype. When studying the effect of HIF-1 degradation, we keep the mtROS production rate fixed, and *vice versa*.

To analyze the effect of the HIF-1 degradation rate, three values of *k_H_*, representing relatively high, moderate, and low degradation rate of HIF-1, are selected. For each value of *k_H_*, we randomly sampled all other parameters using the aforementioned randomization procedure. Again, we collect all stable state solutions from 1000 sets of parameters, and use HCA and PCA to classify the solutions. We find that increasingly stabilized HIF-1, represented by a lower degradation rate, enables a higher percentage of the hybrid metabolic and glycolytic states and lower percentage of the OXPHOS states (**Fig. 4A**). Similarly, we performed an analogous analysis to analyze the effect of mtROS production rate. We find that high mtROS production rate enables a higher percentage of the hybrid metabolic and OXPHOS states and lower percentage of the glycolytic states (**Fig. 4B**). Interestingly, although both stabilization of HIF-1 and elevated production of mtROS can promote the hybrid metabolic state, their effects on the glycolysis and OXPHOS state are opposite. The results here is indicative of the fact that a hybrid metabolic state is rarely observed in normal cells and this appears to be due both to the unstable HIF-1 and relatively low production rate of mtROS relative to cancer cells. Another interesting observation is the emergence of a cluster characterized by low activities of AMPK and HIF-1 and low rates of glycolysis and OXPHOS, referred to as the ‘low-low’ cluster. This state appears especially when the degradation rate of HIF-1 is high and/or the production rate of mtROS is low (**Fig. 4**). In a later section, a subpopulation of single BC cells with the ‘low-low’ characterization is identified and the significance of such a quiescent metabolic phenotype is discussed.

**Figure 4.**
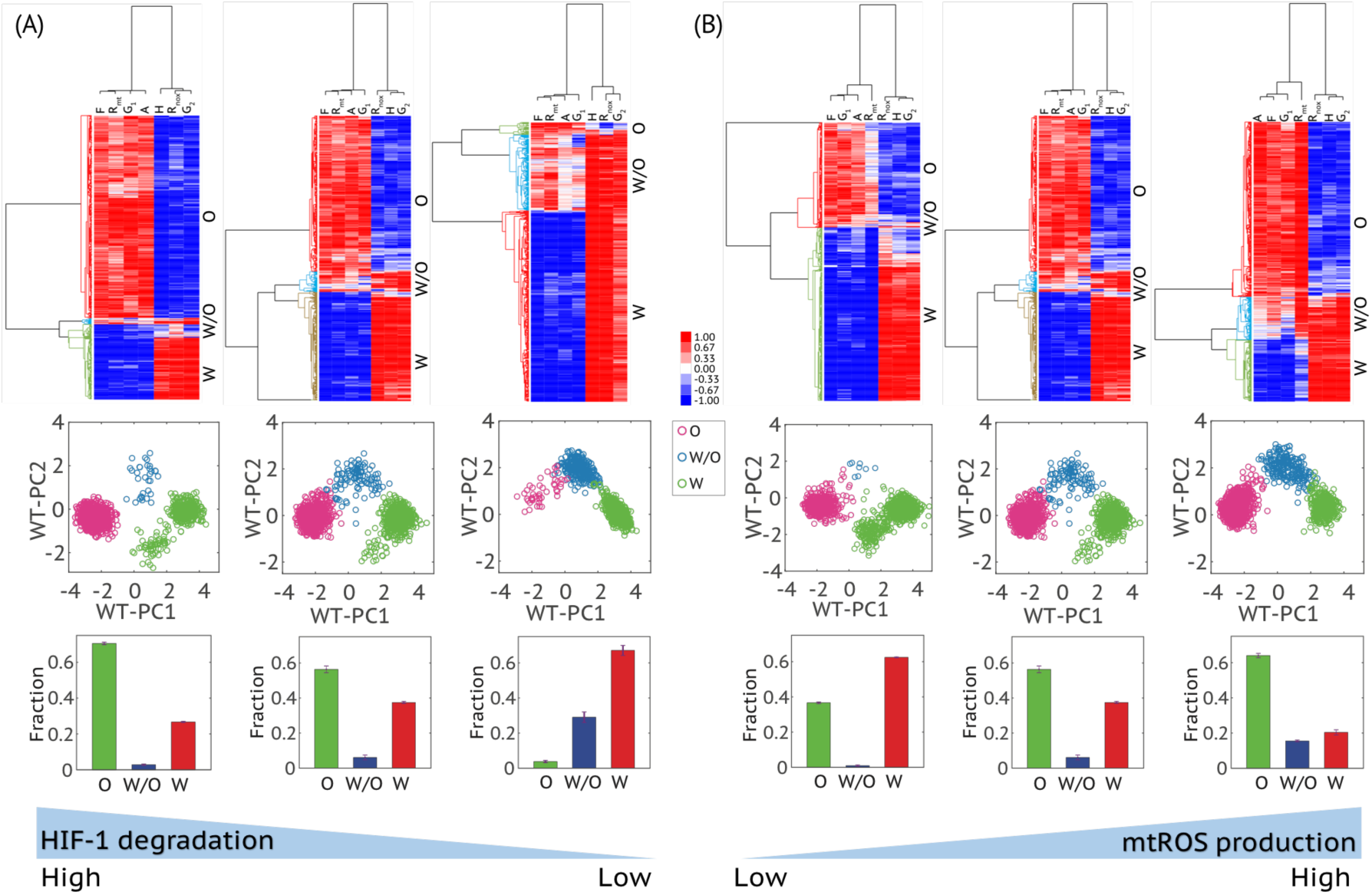
The effects of the HIF-1 degradation rate (A) and the mtROS production rate(B) on metabolic phenotypes. (A) Top panels, hierarchical clustering analyses of the stable state solutions (referred to as samples here) of 1000 sets of parameters with the degradation rate of HIF-1 (*k_H_*) being 0.45 *h*^−1^, 0.25 *h^−1^* and 0.05 *h^−1^* (from left to right). Middle panels, projection of the clustered samples in the top panels onto the PC1 and PC2 generated by the wild type samples with the degradation rate of HIF-1 being 0.25 *h*^−1^ (referred to as WT-PC1 and WT-PC2). Bottom panels, the fractions of the metabolic states - “O”, “W/O” and “W” - corresponding to the top panels. The analysis was repeated for three times and error bars were added. (B) Top panels, hierarchical clustering analyses of the stable state solutions of 1000 sets of parameters with the production rate of mtROS 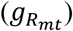 being 30 *μM/min*, 50 *μM/min* and 80 *μM/min* (from left to right). Middle panels: projection of the clustered samples in the corresponding top panels onto the PC1 and PC2 generated by the wild type samples, with the production rate of mtROS being 50 *μM/min*. Bottom panels, the fractions of the metabolic states - “O”, “W/O” and “W” - corresponding to the top panels. The analysis was repeated for three times and error bars were added. The middle figures in (A) and (B) are the same with the parameters *k_H_ = 0.25 h^−^*^1^ and 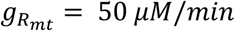, representing the wild type. Z-scores of the stable state solutions were used for clustering analysis and PCA. The solutions of all scenarios here were normalized using the mean and standard deviation of the wild type.

### Evaluating metabolic pathway activity by metabolite abundance

To test the predicted genetic and metabolic characterization of differing cancer metabolism phenotypes, we wish to compare the AMPK/HIF-1 activity and the metabolic pathway activity using metabolomics and transcriptomics data from breast cancer patients’ samples. Note however that the active form of AMPK is its phosphorylated form (pAMPK) and the most important property of HIF-1 is protein stability, neither of these features can be directly captured by the mRNA expression of AMPK and HIF-1. In the previous work, we developed AMPK and HIF-1 signatures to quantify the activity levels of AMPK and HIF-1 by evaluating the expression of their downstream target genes (a total of 33 AMPK downstream genes and 23 HIF-1 downstream genes) (26). The AMPK and HIF-1 signatures were derived by performing PCA on the gene expression data independently for AMPK- and HIF-1-downstream genes, from which the first principal components (PC1s) are used to quantify the activity of AMPK and HIF-1. The AMPK and HIF-1 signatures have been shown to capture the key metabolic features of multiple types of tumor samples from TCGA, such as invasive breast carcinoma, HCC, lung adenocarcinoma (LUAD). Similar findings was also observed in the single cell analysis of LUAD (26). Particularly, a significantly strong anti-correlation between the AMPK activity and the HIF-1 activity has been observed across the aforementioned tumor samples and single cells, where there is no such clear correlation observed between AMPK and HIF-1 gene expression (**Figs. S3–4**).

Here we apply the AMPK and HIF-1 signatures to quantify the AMPK and HIF-1 activity of 45 human BC samples and 45 corresponding adjacent benign breast tissue samples (35) (See **Methods and Materials** for more details). The BC samples show significantly higher HIF-1 activity and lower AMPK activity on average relative to the adjacent benign breast tissues (**Fig. 5A, left panel**), indicating the enhanced glycolytic activity in the BC samples. Moreover, the cancer samples are more stretched in the space of AMPK and HIF-1 signatures with some samples exhibiting very high HIF-1 activity and some samples exhibiting very low HIF-1 activity relative to benign tissues, suggesting heterogeneity in cancer metabolic activities (**Fig. 5A, left panel**). Moreover, a strong anti-correlation between AMPK and HIF-1 activity is identified across the BC samples, which is consistent with our previous results (26).

**Figure 5.**
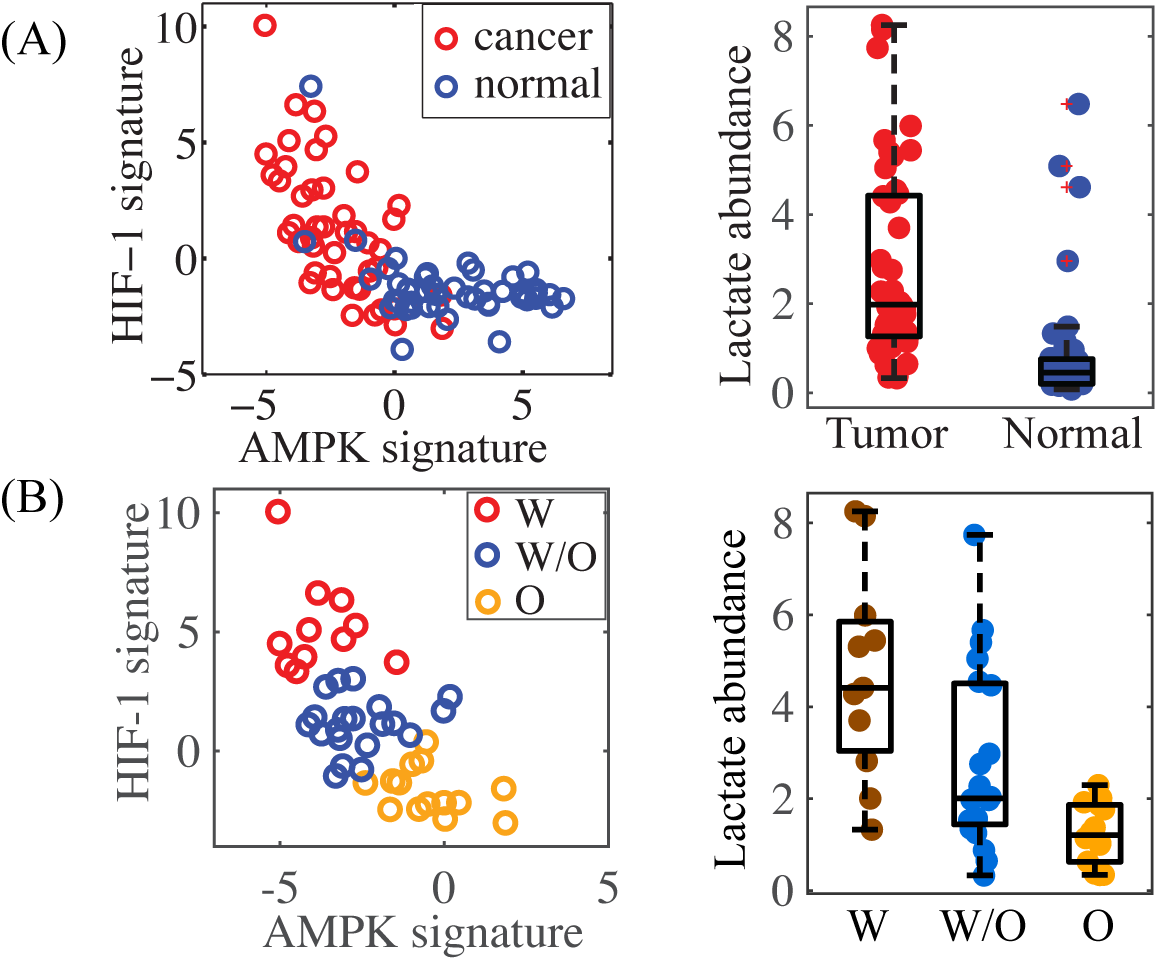
Association of the AMPK/HIF-1 activity with lactate abundance in human breast tumor samples. (A) Left panel: evaluating the AMPK and HIF-1 activities of breast cancer patients’ samples (n=45) and the corresponding normal samples (n=45). Right panel: The relative abundance of lactate in tumor and normal samples (P value < 0.0001). (B) Left panel: Strong anti-correlation of AMPK and HIF-1 activity of tumor samples (Pearson correlation, r = −0.67, P value < 0.0001). The standard k-means clustering analysis was applied to group the tumor samples into the “W”, “O” and “W/O” states. Right panel: The relative abundance of lactate of samples in the “W”, “O” and “W/O” states (P_W-W/O_ < 0.05, P_W/O-O_ < 0.01).

Our new model directly connects these regulatory activities to actual metabolic flux. To test our approach, we need to characterize metabolic pathway activity. We first consider using metabolite abundance. HCA were performed on BC samples and/or benign tissue samples based on the abundance of major metabolites involved in glycolysis, TCA and FAO (**Fig. S5**). The full list of metabolites used here can be found in **Table S1**. The clustering result shows that relative to the benign tissues, BC samples have higher abundance of most metabolites (**Fig. S5**) and among BC samples, the samples exhibit either high abundance of most metabolites or low levels of almost all metabolites (**Fig. S6**). There are no signs of specific metabolic states. These results suggest that using the static metabolite abundance may not be informative in evaluating the metabolic pathway activity. Since most of the metabolites are pathway intermediates, their abundance may not be indicative of the pathway activity since both active metabolic flux (active production) or stalled metabolic flux can cause the accumulation of an intermediate. However, if the metabolite is the end product of a metabolic pathway, its abundance should be indicative of the pathway activity. Since lactate is the end product of glycolysis, we can use its abundance to evaluate the glycolytic activity. The BC samples clearly show higher average abundance of lactate relative to the benign tissues (**Fig. 5A, right panel**). This indicates higher glycolytic activity in cancer tissues and is consistent with the predictions by the AMPK and HIF-1 signatures.

The 45 BC tissues are further classified into three groups based on their AMPK and HIF-1 activities using k-means clustering (**Fig. 5B, left panel**). Among the three groups, tumor samples characterized with highest HIF-1 activity, labelled as ‘W’, exhibit highest abundance of lactate and tumor samples characterized with highest AMPK activity, labelled as ‘O’ exhibit lowest abundance of lactate and tumor samples characterized with both AMPK and HIF-1 activities, labelled as ‘W/O’ show intermediate levels of lactate (**Fig. 5B, right panel**).

Motivated by the ability of lactate abundance to distinguish different metabolic states, we also searched for other metabolites that exhibit significantly differential levels between BC samples and benign tissue samples. Such metabolites include phosphoethanolamine, glutamate, fumarate and cystine in addition to lactate (see the volcano plots in **Fig. S7**). Phosphoethanolamine shows the most differential abundance between BC and benign tissues (*log_2_* tumor/normal = 11.16) and its enrichment in BC has been frequently observed while poorly studied (36, 37). Glutamate enrichment that is often due to the overexpression of glutaminase is a characteristic of breast cancer (38). Accumulation of fumarate, referred to as an oncometabolite, has been shown to trigger epithelial-mesenchymal transition (EMT) in renal cancer (39). Last but not least, cystine enrichment in BC samples identified here is reminiscent of the result that cystine deprivation can induce rapid programmed necrosis in the basal-like BC cells (40). We also perform an analogous analysis to identify the significantly differentially enriched metabolites among groups ‘W’, ‘WO’ and ‘O’ and such metabolites include glutamate and lactate. The list of the significantly differentially enriched metabolites are provided in **Tables S2–4**. All told, however, there is no simple way to accurately estimate metabolic pathway activity directly from metabolite abundance.

### Evaluating metabolic pathway activity by enzyme gene expression

To more directly evaluate the metabolic pathway activities, we develop a new metabolic pathway scoring metric by evaluating the gene expression of key enzymes involved in specific metabolic pathways. The assumption here is that higher metabolic pathway activity would require higher levels of enzymes functioning in that pathway. With this assumption, a total of 14 enzyme genes of FAO, 10 enzyme genes of TCA, 8 enzyme genes of glycolysis were selected as the signature genes to evaluate the activities of OXPHOS and glycolysis. The full list of all enzyme genes used here can be found in **Materials and Methods**. To unbiasedly test for the association of the AMPK and HIF-1 activities with the metabolic pathway activities, we ensured that the genes used in constructing the AMPK and HIF-1 signatures do not overlap with the genes comprising the metabolic pathway scoring metric. The metabolic pathway scoring metric is defined as the average gene expression of relevant enzymes for each pathway. A detailed formulation of this metric can be found in **Materials and Methods**.

First, we use the metabolic pathway scoring metric to quantify the pathway activities of the 45 BC patients’ samples. We perform HCA on microarray data for the enzyme genes and classify the samples into three groups with the most significant gene expression patterns; group ‘W’ contains samples that have high expression of glycolytic enzyme genes and low expression of OXPHOS enzyme genes, group ‘O’ contains samples that have high expression of OXPHOS enzyme genes and low expression of glycolytic enzyme genes and group ‘W/O’ contains samples that have high expression of both OXPHOS and glycolytic enzyme genes, indicating the use of both metabolism phenotypes by the tumor (**Fig. 6A, left**). Then we calculate the pathway scores for each group. The glycolysis score shows a significantly decrease from group ‘W’, to group ‘W/O’ to group ‘O’ and the FAO score of group “O” and “W/O” is significantly higher than that of group “W” (**Fig. 6A, right**). We then evaluate the activity of AMPK and HIF-1 of each sample by the AMPK and HIF-1 signatures. Strikingly, members of group ‘W’ characterized by high expression of glycolytic enzymes show high HIF-1 activity and members of group ‘O’ characterized by high expression of TCA and FAO enzymes show high AMPK activity and finally members of group ‘W/O’ characterized by high expression of both glycolysis and OXPHOS enzyme genes show both AMPK and HIF-1 activities (**Fig. 6A, right**). Importantly, though there are only 45 samples analyzed, a distinct gene expression pattern of enzymes are clearly observed and are strongly associated with the AMPK/HIF-1 activities, as we predicted using our model.

**Figure 6.**
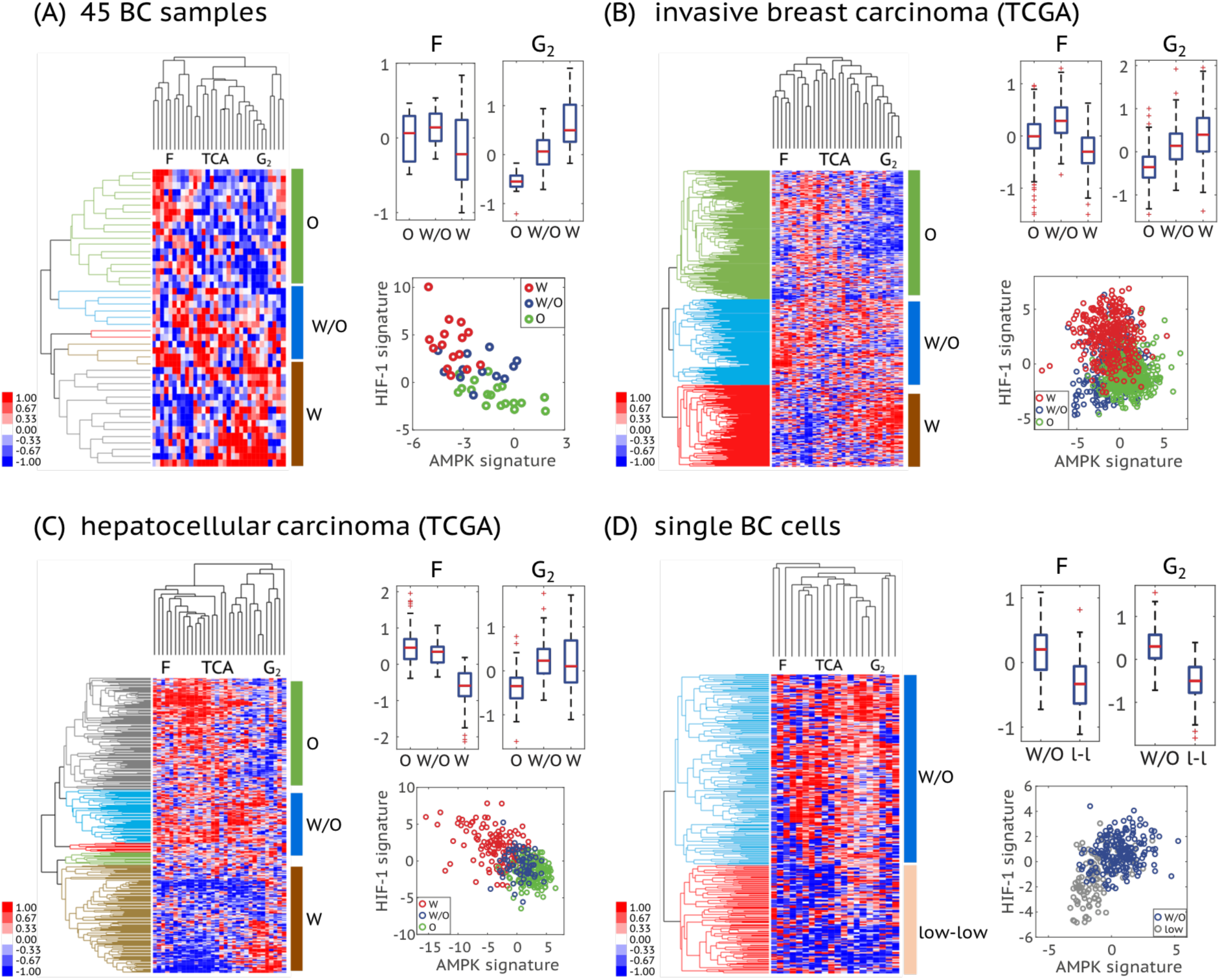
Association of the AMPK/HIF-1 activity with the metabolic pathway activity in 45 BC samples. (A), 1100 invasive breast carcinoma samples from TCGA (B), 373 HCC samples from TCGA (C) and 317 single BC cells (D). In (A) - (C), left panel, average linkage hierarchical clustering analysis of enzyme gene expression (microarray data for (A) and RNA-seq data for (B), (C)) of patient samples. The similarity metric used here is based on Pearson correlation. Each row represents a patient sample and each column represents the expression of one enzyme gene. Three major enzyme gene states - “O”, “W/O” and “W” - are identified and highlighted by different colors in the dendrogram. The cutoff values to get these clusters are 0.04 for (A), 0.1 for (B) and 0.06 for (C). Right panel (top), box plots for the FAO (F) and glycolysis (G_2_) scores of the clustered patient samples. For 45 BC samples, FAO score, *P_O-WO_ = 0.28*, *P_W-WO_ = 0.12*, *P_O-W_ = 0.32*; glycolysis score, *P_O-WO_ <* 0.0001, *PW-WO <* 0.001, *P_o-w_ <* 0.0001; For invasive breast carcinoma, FAO score, *p_O-WO_ < 0.0001*, *p_W-WO_ < 0.0001*, *p_O-W_ <* 0.0001; glycolysis score, *p_O-WO_ <* 0.0001, *p_W-WO_ <* 0.0001, *p_O-W_ <* 0.0001; For HCC, FAO score, *P_O-WO_ <* 0.01, *P_W-WO_ <* 0.0001, *P_O-W_ <* 0.0001; glycolysis score, *P_O-WO_ <* 0.0001, *P_W-WO_ = 0.34, p_O-W_ <* 0.0001; Right panel (bottom), the AMPK and HIF-1 signatures of the clustered patient samples. Here, each hollow dot represents one patient sample. In (D), left panel, average linkage hierarchical clustering analysis of the RNA-seq data of enzyme genes of single BC cells. The enzyme genes whose expression was detected in more than 150 single cells (~half of the total) were used to do clustering. The similarity metric used here is based on Pearson correlation. Each row represents a single cell and each column represents the expression of one enzyme gene. Two major enzyme gene states – “W/O” and “low-low” (low expression of almost all enzyme genes studied here) – are identified and highlighted by different colors in the dendrogram. The cutoff value to get these two clusters is 0.009. Right panel (top), box plots for the FAO (F) and glycolysis (G_2_) scores of cells in the “W/O” and “low-low” clusters. FAO score, *p* < 0.0001; glycolysis score, *p* < 0.0001. Right panel (bottom), the AMPK and HIF-1 signatures of the single cells in the ‘W/O” cluster and the “low-low” cluster. For the heatmaps in (A) - (D), the names of the columns are listed as the names of the metabolism pathways and the full list of enzyme genes can be found in the **Table S8**. The TCA score of each case is shown in **Fig. S8**. For all box plots here, p-values for a balanced one-way ANOVA are calculated.

We further extended the analysis of pathway activity and AMPK/HIF-1 activities using patient data from TCGA. We used HCA to classify 1100 invasive breast carcinoma samples (**Fig. 6B, left**) and 373 HCC samples (**Fig. 6C, left**) into three groups respectively based on the aforementioned enzyme gene expression patterns and calculated their pathway scores. Consistent with what we found in the 45 BC samples, TCGA analysis also allows for the identification of three groups (‘W’, ‘O’ and ‘W/O’). Since we have data from relatively large number of samples, the three metabolic groups of invasive breast carcinoma and HCC samples exhibit more significant differences especially among the FAO and glycolysis scores (**Fig. 6B, C, right**). As predicted, invasive breast carcinoma and HCC samples with high glycolysis score showed high HIF-1 activity and HCC samples with high FAO score showed high AMPK activity and HCC samples with high/intermediate scores of FAO and glycolysis exhibited both AMPK and HIF-1 activities (**Fig. 6B, C, right**). All these confirm the modeling predicted association of the AMPK/HIF-1 activity with pathway activities.

We further validated the association of the AMPK/HIF-1 activity with pathway activity at the single cell level. We looked into the single-cell RNA-Seq data of 317 BC cells (41), which are classified into a ‘W/O’ cluster and a ‘low-low’ cluster (**Fig. 6D, left**), containing cells having high and low expression of both glycolytic and OXPHOS enzyme genes, respectively. Cells in the ‘W/O’ cluster exhibit significantly higher FAO, TCA and glycolysis scores as compared to cells in the ‘low-low’ cluster (**Fig. 6D, right, Figure S8D**). As predicted, cells in the ‘W/O’ cluster exhibit both high AMPK and high HIF-1 activity and cells in the ‘low-low’ cluster exhibit low activity of both (**Fig. 6D, right**). To compare the metabolic state of these BC single cells with benign breast tissues, we projected these BC cells to the AMPK/HIF-1 axes generated by the 45 BC samples and 45 adjacent benign tissue samples (**Fig. S9**). A consistent AMPK/HIF-1 activity characterization is observed for cells in the ‘W/O’ cluster and ‘low-low’ cluster. Intriguingly, some cells in the ‘low-low’ cluster exhibit even lower HIF-1 activity relative to benign tissues (**Fig. 6D, right**). These results confirm the association of the AMPK/HIF-1 activity with pathway activity at the single cell level and also demonstrate the presence of single cell hybrids.

In summary, though the AMPK and HIF-1 activity and the metabolic pathway activity were evaluated independently using different sets of genes, the strong correlation observed at both the tumor level and the single cell level supports the prediction of the model regarding the genetic and metabolic characterization of each metabolic phenotype.

### Experimental validation of the coupling of AMPK/HIF-1 activity with metabolic pathway activity in metastatic cancer cells

To further validate the predictions regarding the coupling of gene activity and metabolic pathway activity in cancer cells, we used various metabolic inhibitors to perturb the glycolysis and/or FAO activity and analyzed the change in AMPK and HIF-1 activity in BC cells. First, we show that the metastatic TNBC cells MDA-MB-231 and SUM-159-PT, that have a significant dependency on mitochondrial FAO (12), exhibit a hybrid metabolic phenotype with both high OXPHOS and high glycolysis, as measured by seahorse respiration analysis (**Fig. 7A, B**). When these TNBC cells were treated with a FAO inhibitor etomoxir (ETX), which prevents the entry of fatty acid into mitochondria by inhibiting the FAO rate-limiting enzyme CPT1, oxygen consumption rate (OCR) was sharply decreased in both MDA-MB-231 and SUM-159-PT cells (**Fig. 7A, B**). Interestingly, ETX-mediated reduction in respiration resulted in a simultaneous increase in glycolysis represented by extra cellular acidification rate (ECAR). This confirmed the metabolic plasticity of these cells. By comparison, only limited ETX-mediated metabolic changes have been observed in an estrogen receptor positive (ER+) BC cell line MCF-7 due to its minimal dependency on FAO (**Fig. 7A**).

**Figure 7:**
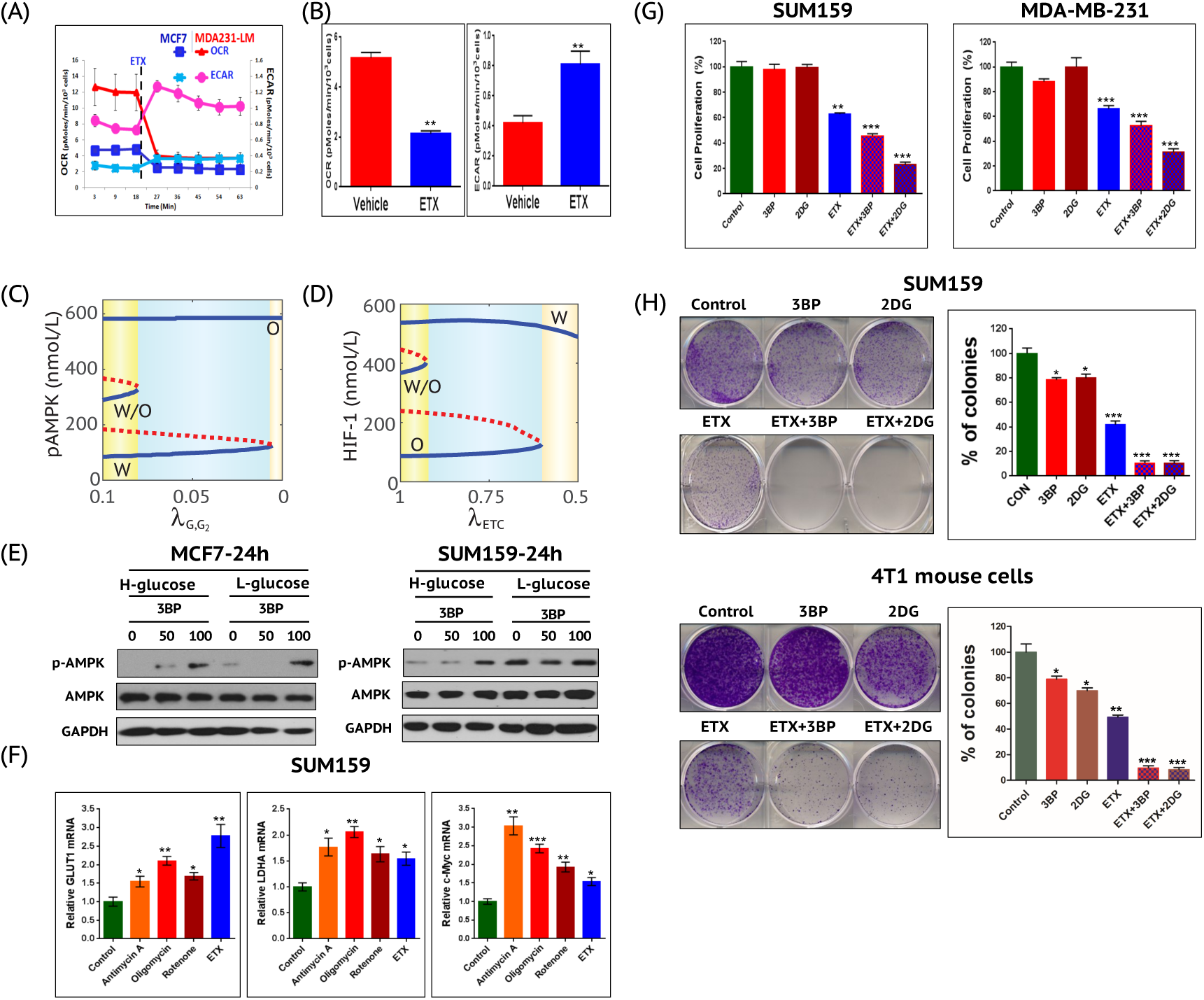
TNBC cells in hybrid metabolic phenotype exhibit maximum clonogenicity relative to cells treated with glycolytic and/or OXPHOS inhibitors. Seahorse XF analysis suggesting that the TNBC cells MDA-MB-231 (A) and SUM159-PT (B) exhibit a hybrid metabolic phenotype and addition of FAO inhibitor (ETX) decreases the respiration and increases glycolysis. (C) Bifurcation diagram of pAMPK levels in response to inhibition of the glycolytic pathway (G2). 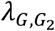 represents the strength of inhibition of G2. The smaller 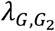, the stronger the inhibition. (D) Bifurcation diagram of HIF-1 levels in response to inhibition of ETC. λ*_ETC_* represents the strength of inhibition of ETC. The smaller λ*_ETC_*, the stronger the inhibition. More details to calculate (D) can be found in **SI section 3**. (E) At 24h following culture of high glucose (4.5 g/L) or low glucose (1 g/L), cells were treated with 50 uM or 100 uM of 3BP for another 24h. The expression and phosphorylation level of AMPK were determined by Western Blotting analysis. GAPDH was used as a loading control. (F) Cells were treated with the mitochondrial ETC complex I inhibitor Rotenone (10 nM), the complex III inhibitor Antimycin-A (10 uM), the complex V inhibitor Oligomycin (5 ug/ml), or ETX (100 uM) for 24h. Expression of glycolytic genes (GLUT1, LDHA, c-Myc) was determined by quantitative PCR. Glycolytic and OXPHOS inhibitors inhibited the cell proliferation (G) and the colony formation (H) in SUM159-PT and mouse metastatic basal breast cancer 4T1 cells. Use of both glycolytic and OXPHOS inhibitors almost eliminates the colony formation potential of SUM159-PT and 4T1 cells. All Data are presented as the means ± standard deviations (SDs). ‘*’ represents P value < 0.05 (two-tailed t-test). ‘**’ represents P value < 0.01. ‘***’ represents P value < 0.001.

To analyze how altering metabolic pathway activity affects the gene activity, we first use the model (**Fig. 2**) to simulate how repressing glycolytic pathway affects AMPK activity. Increasing inhibition of glycolysis first excludes the existence of the hybrid metabolic state, then the glycolytic state and finally only the OXPHOS state remains (**Fig. 7C**). In other words, the cell populations are expected to be more OXPHOS with increased average pAMPK levels upon glycolytic inhibition. We also analyze how repressing ETC affects HIF-1 activity. Increasing inhibition of ETC first excludes the existence of the hybrid state, then the OXPHOS state and finally only the glycolysis state remains (**Fig. 7D**). To test the model prediction, we used a glycolytic inhibitor, 3-bromopyruvate (3BP), to treat the BC cells and we found that AMPK was activated, i.e. the levels of pAMPK are upregulated in both MCF-7 cells and SUM-159-PT cells within 24h, especially in high glucose conditions (**Fig. 7E**). When the mitochondrial function of SUM-159-PT cells is repressed, expression of the glycolysis relevant genes, including GLUT1, LDHA and c-Myc was significantly upregulated, indicating the increased glycolytic activity at the population level (**Fig. 7F**). These experimental results support the model prediction.

To further understand the functional significance of various metabolism phenotypes in the tumor properties, we performed proliferation assays and clonogenic assays using SUM-159-PT and mouse metastatic basal breast cancer 4T1 cells. While glycolytic inhibitors 3BP and 2-deoxyglucose (2DG) showed minor inhibition in their clonogenic potential but not in proliferation, their combination with ETX caused a major reduction in both the proliferation and clonogenic potential of both of the cell lines (**Fig. 7G, H**). This suggests that disrupting the survival potential of either tumors or individual cells with hybrid metabolic status requires targeting both metabolic pathways.

## Discussion

For a long time, aerobic glycolysis has been regarded as the dominant metabolic phenotype in cancer. However, there has recently been increasing experimental work demonstrating a critical role of oxidative phosphorylation and mitochondrial biogenesis in tumorigenesis and metastasis. Apparently, cancer cells are able to adjust their metabolism phenotypes to adapt to the microenvironment. In this regard, we showed that cancer cells can acquire a stable hybrid metabolic state by a demand-sensitive crosstalk of regulatory proteins and energy pathways (26).

In this study, we established a theoretical framework (**Fig. 2**) that couples the gene regulatory circuit with metabolic pathways in order to explore the genetic and metabolic interplay between glycolysis and OXPHOS. The model predicts a direct association of high AMPK activity with high OXPHOS activity and high HIF-1 activity with high glycolysis activity. To validate this prediction, we developed signatures to quantify the activity of metabolic pathways and could therefore show the association of the pathway activity with the AMPK and HIF-1 activities, evaluated by our previously defined AMPK and HIF-1 signatures. By applying the pathway signatures and the AMPK/HIF-1 signatures, we confirmed their association and the existence of the hybrid metabolic phenotype at both the tumor level and the single cell level. Furthermore, we performed experiments in TNBC using MDA-MB-231 and SUM-159-PT cells, which stably maintain a hybrid metabolic phenotype. Inhibiting the glycolytic activity in these TNBC cells activates AMPK. With the use of selective inhibitors, we show that cells in the hybrid metabolic phenotype exhibit maximum proliferation and clonogenicity relative to cells in a more glycolytic or more OXPHOS phenotype. To the best of our knowledge, our work is the first to couple the gene regulation with the metabolic regulation to elucidate cancer metabolic plasticity, through integrating modeling, data analysis and experiments.

The hybrid metabolic phenotype, characterized by high HIF-1/AMPK activities and high glycolysis/OXPHOS (glucose oxidation and FAO) activities, enables for tumors and even for individual cancer cells the metabolic plasticity to utilize various kinds of nutrients, such as glucose and fatty acid. It allows the cells to efficiently produce energy through multiple metabolism pathways and meanwhile synthesize biomass for rapid proliferation using byproducts from glycolysis. In addition, the hybrid metabolic phenotype maintains the cellular ROS at a moderate level so that cancer cells can benefit from ROS signaling (42) and still avoid DNA damage due to excessive ROS (43). Moreover, the hybrid metabolic phenotype may be specifically associated with metastasis (10, 12, 13, 16, 17) and the survival and propagation of therapy-resistance cancer stem cells (CSCs) [38–40]. As we showed in **Fig. 7**, combination of the glycolytic and OXPHOS inhibitors, effectively eliminates the tumor survival potential of hybrid cells. Dual inhibition of glycolysis (by 2-DG) and OXPHOS (by metformin) has been shown to effectively repress tumor growth and metastasis across multiple preclinical cancer models (44). Moreover, the hybrid metabolic phenotype has recently been observed in immune cells. The regulatory T cells using both glycolysis and FAO exhibit more successful expansion than the conventional T cells using primarily glycolysis (45). Future work should develop in more detail the nature of the hybrid metabolic phenotype and its coupling with other hallmarks of cancer, such as metastasis and stem-like properties. Another intriguing result is the emergence of a ‘low-low’ metabolic state with the characteristic feature of low AMPK/HIF-1 activity and low OXPHOS/glycolysis activity, especially when the HIF-1 degradation rate is high or mtROS production rate is low, as predicted by the model (**Fig. 4**) and shown by gene expression data analysis (**Fig. 6**). This metabolically inactive state has been observed in bacterial persister cells (46) and its significance in cancer is worthy of future study.

In this work, we focused on the genetic and metabolic interplay of glycolysis and OXPHOS (FAO and glucose oxidation). It is worth noting that glutamine oxidation, that is often driven by the oncogene MYC, can also play a critical role in regulating tumor growth and metastasis (47, 48). Moreover, abnormal metabolism, as a hallmark of cancer, involves not only cancer cells. The surrounding glycolytic cancer-associated fibroblasts (CAFs), the stromal cells which often dominate the tumor microenvironment, can provide energy-rich metabolites to promote OXPHOS activity and anabolic metabolism of cancer cells (49). These multicellular aspects, although beyond the scope of this work, are definitely worthy of further investigation combining both theoretical and experimental efforts.

## Materials and Methods

### 1. Formulation of the mathematical model

The generic deterministic equations representing the temporal dynamics of pAMPK, HIF-1, mtROS, noxROS are given by,

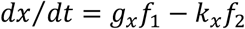

where *g_x_* and *k_x_* represent the basal production and degradation rates of component *x*, *f*_1_ and *f*_2_ are two functions representing the regulation of *x*’s production and degradation due to the cross-talk between *x* and other components. Since the chemical reactions in metabolism processes are much faster than the gene regulation, we assume the metabolite concentrations and the metabolic pathways are in the equilibrium state at certain level of pAMPK and HIF-1. The uptake of glucose is regulated by AMPK and HIF-1. The intracellular glucose is shared by glucose oxidation and glycolysis. The production of acetyl-CoA is from glucose oxidation and fatty acid oxidation. The generation of ATP takes place from all three metabolic pathways studied here. The detailed modeling procedure can be found in **SI section 1, 2** and the parameter values can be found in **Table S5**.

### 2. The metabolic pathway scoring metric

The metabolic pathway score is defined as,

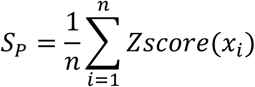

Where *x_i_* represents the expression of enzyme gene *i* and *n* represents the total number of genes analyzed for pathway *P*.

### 3. 45 BC samples and 45 benign tissue samples

67 human BC samples and corresponding adjacent benign breast tissue samples are provided in (35). The microarray data and metabolite data of 45 BC samples and their paired adjacent benign tissue samples are selected for analysis in the present study. For other paired BC and benign samples, the AMPK and HIF-1 downstream gene expression of either BC or benign sample is missing. The sample ID used in this study is listed in **Table S6**.

### 4. Experimental design

#### Cell Culture and Chemicals

Human breast cancer cell line MDA-MB-231 and MCF-7 cells were from the American Type Culture Collection (ATCC), and SUM-159-PT cells from Asterand Bioscience was provided by Dr. Dave (MD Anderson Cancer Center). Cells were cultured in Dulbecco’s Modified Eagle’s Medium (DMEM) supplemented with 100 μg/mL of streptomycin, 100U/mL of penicillin and 10% heat-inactivated fetal bovine serum (FBS) (Genedepot). Etomoxir (ETX; Tocris), 2-deoxyglucose (2-DG; Sigma) and 3-bromopyruvate (3-BP; Sigma) were dissolved according to manufacturer’s instruction.

#### Cell Respiratory Assay

Oxygen consumption rates (OCR) and extracellular acidification rate (ECAR) were measured using the mitochondrial stress test procedure in XF media (non-buffered DMEM containing 5 mM glucose and 2 mM sodium pyruvate; note lack of buffer is absolutely required to measure pH drop that indicates ECAR) with the XF96 Extracellular Flux Analyzer (Seahorse Bioscience) (50).

#### Cell Proliferation Assay

The sulforhodamine B (SRB) based colorimetric assay was also conducted to rule out artifacts from mitochondrial alterations in MTT activity. Briefly, cells were fixed with 5% trichloroacetic acid to terminate reaction, and 0.4% SRB (Sigma) in 1% acetic acid was added to each well. After 30 min incubation, the plates were washed with 1% acetic acid, and dyes were dissolved by 10 mM Tris buffer. Then, the absorbance density values were read by Infivite M200 PRO reader (510 nm). Experiments were performed in triplicates.

#### Clonogenic Assay

Cells (1×10^3^/well) were seeded into each well of a six-well plate, with three technical replicates per condition. Drug-containing media were changed every 3 days for 2 weeks. Cells were rinsed with PBS, fixed in fixation solution (10% methanol, 10% acetic acid, 80% H_2_O) for 10 min and stained with 0.5% crystal violet (20% methanol, 80% H_2_O) for 30 minutes. Crystal violet was solubilized with 10% acetic acid for 15 min and quantified by absorbance at 590 nm.

#### RNA Isolation and Quantitative Polymerase Chain Reaction

Total RNA was isolated using mRNeasy extraction kit (Qiagen). To quantify gene expression, RNA samples were submitted to reverse transcription and real-time PCR using primers listed in **Table S7**. cDNA was amplified using the MX3000P qPCR System (Stratagene) and SYBR Green Supermix (Bio-Rad). The relative quantification of mRNA expression was calculated using the 2^−ΔΔCt^ method.

#### Western Blotting

Cells were washed with ice-cold PBS and cell lysates were prepared in RIPA buffer containing phosphatase inhibitors and protease inhibitors (Gendepot). Cell lysates were centrifuged at 14,000 rpm for 15 min at 4°C, and supernatants were collected. Equal amounts of protein were separated by SDS-PAGE gel and transferred onto nitrocellulose membranes (Bio-Rad). The membrane was blocked with a 5% skim milk solution and incubated with the following antibodies: anti-phospho-AMPK, anti-AMPK and anti-GAPDH (Cell Signaling). The ClarityTM western ECL substrate kit (Bio-Rad) was used to detect bound antibodies. The signal was detected by x-ray film.

## Acknowledgements

D.J., J.N.O and H.L. are supported by the Physics Frontiers Center National Science Foundation (NSF) grant PHY-1427654 and the NSF grant PHY-1605817. M.L. is supported by the start-up fund from the Jackson Laboratory, the National Cancer Institute (NCI) grant P30CA034196, and the National Institute of General Medical Sciences grant R35GM128717. J.H.P., K.H.J. and B.A.K. are supported by the NCI grants R21CA173150, R21CA179720 and R03CA212816, Collaborative Faculty Research Investment Program (CFRIP), Breast Cancer Research Foundation (BCRF), Dan L Duncan Comprehensive Cancer Center (DLDCC) and Charles A. Sammons Cancer Center. We acknowledge the contribution of Dr. Stefan Ambs from NCI for the data of breast cancer and benign tissue samples.

